# PVR and Nectin-2 blockade trigger macrophage anti-tumor functions, promote immune cell recruitment and prevent cervical tumor growth

**DOI:** 10.64898/2026.01.30.702794

**Authors:** O.M. Diallo, N. Boucherit, T. Fernez, A.O. Barry, E. Billion, M.S. Rouviere, A. Ben Amara, L. Gayraud, J.A. Nunes, X. Carcopino, E. Lambaudie, R. Sabatier, M. Richaud, M. Lopez, A.S. Chretien, M.S. Diallo, D. Olive, L. Gorvel

## Abstract

Despite vaccination, cervical tumors remain a health issue and require treatment improvement. In 2023, Pembrolizumab, an anti-PD-1 immune checkpoint blockade (ICB), was introduced in the case of advanced or metastatic cervical tumors. This treatment significantly increased progression free survival from 20% to 50%. However, some patients remain resistant to anti-PD-1 treatment, which calls for new targets. In our study, we highlighted poliovirus receptor (PVR) and Nectin-2 as potential ICB targets. Indeed, PVR and Nectin-2 are TIGIT ligands, an immunomodulatory checkpoint expressed by regulatory T cells or exhausted T cells. The binding of PVR or Nectin-2 to TIGIT maintains the immunosuppressive signal in the immune cells allowing tumor progression. Furthermore, we observed that PVR and Nectin-2 were highly expressed on tumor cells and tumor associated macrophages (TAMs) accross the different histological subsets of cervical tumors. Therefore, we hypothesized that using anti-PVR and anti-Nectin-2 anti-bodies would lift immunosuppression in cervical tumors. To that end, we used CyTOF to asses precise immunophenotyping of the targets *ex vivo*, high throughput confocal microscopy to assess phagocytosis, monocyte derived macrophages (MDM) coupled with 3D cell culture models to assess the impact of our treatment on MDM and TAM repolarization and tumor growth. We could demonstrate that our treatment repolarized macrophages towards an inflammatory profile and that this was followed by a reactivation of macrophage cytotoxic function such as phagocytosis. We also demonstrated that anti-PVR and Nectin-2 treatment allowed the control of tumor growth in 2D and 3D cell culture models. We could also develop a pre-clinical model of autologous cell culture from cervical cancer patients. Using the MIVO technology, which combine organotypic culture and fluidics, we could assess peripheral blood mononuclear cells recruitment towards tumor cells in the presence or absence of anti-PVR and anti-Nectin-2. In conclusion we could demonstrate that targeting macrophages via the PVR/Nectin-2 couple reactivates cervical tumor growth control and improves immune cell recruitment.

## Introduction

Immune checkpoint blockade (ICB) revolutionized the treatment of cancer, including solid tumors. In cervical cancer, anti-PD-1 immunotherapy (pembrolizumab), was accepted as the standard of care for advanced tumor in 2023 (1). While the results showed and improved survival for 40-50% of patients, the remaining relapsed. This is why the research for new ICB targets in cervical tumors is critical. In that context, the TIGIT-PVRL axis represents a good opportunity to enhance immune response in solid tumors. This axis is complex and involves lymphoid cells, which express TIGIT, CD96 and DNAM-1, tumor cells, which may express the poliovirus receptor (PVR), Nectin-1, Nectin-2 and Nectin-4, and the monocyte/macrophage compartment which may express PVR and Nectin-2. The TIGIT-PVRL axis finely regulates the balance between anti-tumor inflammatory response (PVRL binding to DNAM-1) and pro-tumor immunosuppressive response (PVRL binding to TIGIT)(2–4). In the tumor microenvironment this interaction is forced towards the immunosuppressive PVRL-TIGIT binding, which leads to CD8+ T cell exhaustion and myeloid compartment (dendritic cells and macrophages) polarization towards anti-inflammatory profiles (2,5). The majority of studies in which the TIGIT-PVRL axis was targeted relied on TIGIT blockade to avoid CD8+ T cells exhaustion and reinvigorate them (6,7). In the Skyscraper trial, anti-TIGIT Tiragolumab (IgG4) was used in combination with anti-PDL1 (Atezolizumab) to enhance antitumor response. The results showed an improvement of response to treatment and overall patient survival compared to Atezolizumab alone (8). The Cityscape trial used anti-TIGIT mAbs were used in combination with anti-PD-1 demonstrating a moderate improvement in patient response compared to anti-PD-1 alone. This combination aiming at removing 2 of the CD8^+^ T cell cytotoxic brake (PD-1 and TIGIT) did not meet the expected improvements. To understand this phenomenon, a study was built based on the cityscape results and showed that the responders had a stronger involvement of the myeloid cell compartment. Furthermore, this could be enhanced experimentally by switching the isotype of the antibody from IgG4 to IgG1 or IgG2b in murine models (9). Here, the authors focused on the antibody-dependent cellular phagocytosis (ADCP) function of macrophages, highlighting the importance of macrophage function in the tumor microenvironment (TME). Tumor associated macrophages (TAMs) are one of the most represented cell type in the TME and exhibit a panel of functions which participate to immunosuppression and tumor progression (10,11). For instance, the enrichment of TAMs in a CD8^+^ T cell deprived TME is of poor prognosis in cervical tumors (12). Our team could demonstrate a more differentiated immune checkpoint expressing TAM profile (CD40^+^, PD-L1^+^) in better prognosis cervical tumors harboring tertiary lymphoid structures, while producing more IL-6 in poorer prognosis tumors (13). Indeed, immunosuppressive cytokine production (IL-10, TGF-β) and *in situ* tumor antigen presentation (14) contribute to cytotoxic cell suppression and exhaustion. Furthermore, TAMs are able to remodel the extracellular matrix (matrix metallo-proteases, MMP2 and 9), promote tumor growth (endothelial growth factor, EGF) and to promote angiogenesis (vascular endothelial growth factor, VEGF) (15). Furthermore, macrophages in general and TAMs are highly plastic in phenotypes and functions and are able to switch from immunosuppressive and pro-tumor to inflammatory, phagocytic and anti-tumor. This was demonstrated in studies in which ICB *in vitro* or *in vivo* allowed macrophages functional switch towards inflammation (10,11). Plasticity represents the main advantage to target macrophages and TAMs in solid tumors.

Here, we propose to investigate macrophage response to antibodies blocking PVR and Nectin-2 interaction their ligands in cervical tumors. We could show that PVR and Nectin-2 are mainly expressed by TAMs in the TME and that this expression could be modeled by using conventional MCSF induced (Macrophage colony stimulating factor, also known as CSF-1) monocyte-derived macrophages (MDMs). We demonstrate than anti-PVR and anti-Nectin-2 are able to induce a phenotypic switch from M2-like MDM to M1-like MDMs and to restore tumor cell line phagocytosis in vitro. We also show that targeting PVR and Nectin-2 reduces tumor growth in 2-Dimensional (2D) and immunocompetent 3-Dimensional (3D) spheroid cell cultures. Finally, we could model immune cell recruitment by using a 3D cell culture system coupled to millifluidics. Indeed, we cultured Cervical tumors cells while having autologous PBMCs circulating in the fluidic system and tested the effect of our anti-PVR and anti-Nectin-2 mAbs. Our study demonstrates the potential of using PVR and Nectin-2 blockade alone or in combination to reinvigorate macrophages antitumor functions and enhance immunotherapies.

## Methods

### Human tumor samples

Human cervical tumor samples (n = 14) were obtained from patients who were included in the prospective XAC-03 (NCT02875990) and GC-Bio-IPC (NCT01977274) clinical trials conducted from 2013 to 2025. The XAC-03 and GC-Bio-IPC trials were approved by the Institutional Review Board: *Comité d’Orientation Stratégique, Marseille, France* of the Paoli-Calmettes Institute and the Assistance Publique des Hopitaux de Marseille, respectively. Written informed consent was obtained from all patients according to the Declaration of Helsinki. Inclusion criteria for both XAC-03 and GC-Bio-IPC protocols the following: patients who are more than 18 years of age with cervical cancer and who did not receive any treatment. Patients were excluded if pathology returned any diagnosis of nongynecological tumor cancer.

### Cell lines

SiHa and Caski cervical cancer cell lines were purchased from the ATCC (American Type Culture Collection). They were transduced for mKate2 gene to obtain stable long-term fluorescence.

### Anti-PVR and anti-Nectin-2 antibodies

Anti Nectin 2 and PVR mAbs were produced in the laboratory and harbor the following characteristics. Anti nectin-2 R2.477 (IgG1k) binds the regions formed by the two IgC like domains as described previously (Lopez et al. J Virol p.1267, 2000). Anti-PVR PV404 (IgG1k) binds to the region formed by the two IgC-like domains of PVR as described previously (doi/10.1084/jem.20032206). PV404 mAbs also blocks the TIGIT-PVR interaction as well as the DNAM-1-PVR interaction.

### mRNA expression of PVRL family molecules

Transcript expression of PVRL family genes, *PVR*, Nectin-1 (*PVRL1*), Nectin-2 (*PVRL2*), Nectin-3 (*PVRL3*) and Nectin-4 (*PVRL4*) in cervical tumor sample was assessed using the publicly available The Cancer Genome Atlas (TCGA) CESC dataset. Expression was extracted and analyzed using Phantasus (16).

### Cell enrichment analysis

CIBERSORTx (Stanford University) was used for the immune cell enrichment analysis of the cervical squamous cell carcinoma and endocervical adenocarcinoma (CESC, Database of Genotypes and Phenotypes accession number: phs000178) TCGA gene expression value database (n = 309 samples). The individual gene expression file was uploaded to CIBERSORTx (17), and run using both the relative and absolute modes, signature: LM22, 500 permutations, and disabled quantile normalization (recommended for RNA-seq data). Data were then grouped using hierarchical clustering, and CD8+ enriched, TLS enriched, M1/M2 enriched and M0 enriched patients were identified. Results from CIBERSORTx enrichment are available in **Supp. Table 1**.

### Survival analysis

For survival analyses, overall survival (OS) was defined as the time from diagnosis until death. Patients without an event were censored at the time of their last follow-up. Using the TCGA CESC cohort mRNA expression dataset, survival times of CD8^+^ enriched, TLS enriched, M1/M2 enriched and M0 enriched patients were estimated using the Kaplan–Meier method and compared using the log-rank test.

### Single cell RNA sequencing dataset collection filtering and populations identification

Four publicly available scRNA-seq datasets from cervical cancer and healthy tissue were collected from Gene Expression Omnibus databases (GEO, NCBI). Cell ranger files were imported into R studio and the Seurat package was used for filtering, log-normalization of raw expression matrices, and analysis (18). Exclusion of multiplets, dropouts, dead cells, and low-quality cells was performed as follows, we excluded genes expressed in fewer than 5 cells, cells expressing fewer than 200 genes, and cells with more than 10% of mitochondrial gene counts. For all datasets, we subsampled datasets on the *PTPRC* (CD45) transcript expression (>2) to isolate immune cells and samples containing no or too few (n < 30) viable PTPRC^+^ cells were excluded from the analysis. PTPRC+ cells from the 4 datasets were then integrated to identify anchors cells across datasets. Integated datasets were processed as follows: the top 800 highly variable genes and the first 20 components of the principal component analysis (PCA) were used to compute Uniform Manifold Approximation and Projection for Dimension Reduction (UMAP). The perplexity for UMAP was defined automatically based on the number of cells in the dataset. Clusters were identified by the shared nearest neighbors (SNN) method and annotated based on a differential gene expression analysis of each cluster compared to all other clusters. The top 20 differentially expressed gene for each cluster was entered in the ENrichR software for annotation. PVR and NECTIN-2 log-normalized expression was extracted from the datasets and displayed across clusters. PVR and NECTIN-2 top 20 differentially expressed were used to identify markers that were present in the CyTOF mass cytometry panel and validate the presence of these clusters in GC-Bio and Xac03 cohorts.

### Mass Cytometry

Tumor cells were incubated with cisplatin (1 μmol/L) to stain dead cells. Nonspecific epitope binding was blocked with 0.5 mg/mL Human Fc Block (BD Biosciences). Mononuclear cells were stained for extracellular epitopes for 1 hour at 4°C, followed by permeabilization (Foxp3 permeabilization reagent, eBioscience) for 30 minutes at 4°C and intracellular marker staining with the panel described in **Supp. Table 2**. Cells were then washed and labeled overnight with 125 nmol/L iridium intercalator (Fluidigm) in 2% paraformaldehyde (Thermo Fisher Scientific). Finally, cell pellets were suspended in Milli-Q water (Merck Millipore) containing 10% EQ Four Element Calibration Beads (Fluidigm) and filtered through a 35 μm membrane before acquisition on a mass cytometer (HELIOS instrument, Fluidigm). Myeloid cells (CD45^+^CD3^-^CD19^-^CD56^-^CD33^+^) were manually gated and then exported using FlowJo (Becton-Dickinson). Data were arcsinh-transformed with a cofactor of 5 (19). Myeloid cell subpopulations were manually gated to integrate single cell RNAseq clusters to the analysis and projected onto a Pairwise Controlled Manifold Approximation (PaCMAP) (20) using the OMIQ software suite.

### MDM differentiation and polarization

Peripheral blood Mononuclear cells (PBMCs) were isolated by Ficoll gradient from buffy coats obtained from Etablissement Français du Sang (EFS). Monocytes were then isolated from PBMCs using the CD14+ isolation microbeads (Miltenyi biotech.) following the manufacturer instructions. Briefly, PBMCs were incubated for 15 min with anti-CD14 antibodies, washed and incubated with magnetic microbeads for 15 min at 4°C. PBMCs were passed through a MS column (Miltenyi) fixed on a magnet and CD14-cells were allowed to eluate. After 3 washes in MACS buffer (Miltenyi) the column was removed from the magnet and CD14+ monocytes were flushed in a 15ml tube. Finally, monocytes were adhered in a 6 well plate and culture in the presence of RPMI 1640 (Gibco) supplemented with 10% FCS (fetal calf serum) and macrophage-colony stimulating factor (M-CSF, 40ng/ml) for 5 days to generate M2-like monocyte-derived macrophages (MDM2). For M1-like MDM (MDM1) we added 100ng/ml of IFN-γ on the fourth day to polarize MDM towards an inflammatory profile. Cells were then used in experiments.

### Flow Cytometry

Flow cytometry was used for PVR and Nectin-2 expression as well as MDM polarization assays. MDM were differentiated as stated previously and were or were not incubated with 3µg/ml of soluble recombinant TIGIT-fc (TIGIT-fc), in the presence of control IgG1 (10µg/ml), anti-PVR (10µg/ml), anti-Nectin-2 (10µg/ml) or both anti-bodies (Combo). For PVR and Nectin-2 expression, MDM were incubated with Live/dead Aqua (Invitrogen), anti-PVR BV605 (Biolegend) and anti-Nectin-2 PE (Becton-Dickinson). For MDM polarization assay, MDM were stained for Live/dead Aqua (Invitrogen), CD11b PerCP-Cy5.5 (Biolegend), CD14 APC-H7 (Biolegend), HLA-DR BV605 (Biolegend), CD86 BV650 (Biolegend), CD80 PE (Becton-Dickinson), CD83 FITC (Becton-Dickinson), CD163 PE-Dazzle 594 (Biolegend), CD206 BV421 (Biolegend) and CD209 PE-Cy7 (Biolegend) for 30 min in the dark at 4°C. MDM were then washed and acquired on a BD Symphony flow cytometer (Becton-Dickinson).

### Phagocytosis

MDM were differentiated as described and stained using PKH67 (488nm emission fluorescence). They were then cocultured with mKate2-SiHa (594nm emission fluorescence) cell line at a ratio of 1/5 in a dark 96 flat well plate for 30min, 60min, 120min and 240min in the presence of control IgG1 (10µg/ml), anti-PVR (10µg/ml), anti-Nectin-2 (10µg/ml) or both anti-bodies (Combo). Cell were then imaged on a high throuput confocal microscope and 10 fields with 100 cells were acquired. Data were analyzed to detect the presence of SiHa inside MDMs and the amount of 594nm pixels inside the MDM cytosol. Results were quantified and plotted using Graphpad Prism V8.

### Wound healing assay

MDM and SiHA were co-cultured in inserts (Culture-Inserts 2 Well for self-insertion, Clinisciences) composed of two separated chambers, in the presence of TIGIT-fc (3µg/ml) in the presence or absence of control IgG1 (10µg/ml), anti-PVR (10µg/ml), anti-Nectin2 (10µg/ml) or Combo (10µg/ml). Inserts were then removed to let tumor cells migrate towards each other and fill the gap. Gap area was then quantified by video microscopy for 24h. Data were analyzed by using the FijI software (Image J) and plotted using GraphPad Prism version 8.00.

### Spheroid growth assay

Spheroid were formed from Caski cell lines in the presence of MDM (M-CSF), for which PVR and Nectin-2 expression was confirmed by flow cytometry, during 5 days. Spheroids were then co-cultured in the presence of anti-CD3/CD28 activated T cells, which express TIGIT after 3days, and control IgG1 (10µg/ml), anti-PVR (10µg/ml), anti-Nectin-2 (10µg/ml) or both anti-bodies (Combo). Spheroid size and shape was assessed by video-microscopy (Olympus) during 90h. Videos were analyzed using FijI (Image-J) and spheroid size and shape data were extracted and plotted on Graphpad Prism V8.

### 3D culture with millifluidics (MIVO system)

Patient tumor samples from Xac03 and GC-Bio protocols were thawed and rested for 30 min in RPMI 1640 (Gibco) supplemented with 10% FCS at 37°C + 5% CO2. They were then stained for CD45-BV786 (Biolegend) at 4°C for 30 min and cultured into 25% matrigel onto a 8µm transwell filter overnight in the presence of IgG1 (10µg/ml), anti-PVR (10µg/ml), anti-Nectin-2 (10µg/ml), both antibodies (Combo, 10µg/ml) or Atezolizumab (10µg/ml). The filter containing tumor sample culture was then put on the MIVO system single flow chambers. Autologous CD45-APC stained PBMCs were loaded into the millifluidics. Peristaltic pump was then turned on letting PBMCs flow in the system overnight. Transwell filters were then removed, dissociated using Gentle Dissociation Solution (Stem Cell) and stained for spectral cytometry using the panel in **Supp. Table 3**. Filter supernatants were saved for Luminex dosages. Tumor infiltrating cells were investigated by spectral cytometry using CD45-BV786^+^ cells as first gate. Cell migration was assessed by spectral cytometry in the dissociated cell samples from the transwell filter by analyzing CD45-APC^+^ gating.

### Spectral cytometry

Patient tumor samples used in the MIVO experiments were dissociated and stained for spectral cytometry. Antibodies with excitation from UV and Violet lasers were premixed into Brilliant Buffer (Becton-Dickinson), while antibodies coupled to fluorochromes exited by Blue, Yellow/Green and Red lasers were premixed into Monocyte True Stain Buffer (**Supp. Table 3**). ViaDye Red (CyTEK) viability dye was added to the total mixed and cells were stained for 30 min at 4°C. Cells were ten washed and acquired on the Aurora spectral cytometer (CyTEK).

### Statistical analyses

Statistical analyses were generated using GraphPad Prism version 8.00, with data expressed as mean ± SE of the mean. Statistical significance between two groups was calculated using the nonparametric Mann– Whitney test, whereas multiple group comparisons were calculated using the nonparametric Kruskal– Wallis test, followed by Dunn’s multiple comparison posttest. A P value < 0.05 was considered as significant. Statistical significance of multiple parameters across multiple groups was assessed by using two-way ANOVA with multiple comparisons and FDR assessment (significance: q value < 0.05, with individual P value < 0.05).

## Results

### TAMs are of poor prognosis in cervical tumors

First, we wondered what was macrophages contribution to patient survival n cervical tumors. For that, we used the TCGA CESC dataset and performed a cell enrichment analysis using the CibersortX algorithm and the LM22 signature. The obtained enrichment values (**Supp. Table 1**) were plotted against patients, grouped using unsupervised hierarchical clustering and displayed as a heatmap of all cell types (**Figure 1, A**). Significantly enriched cell type analysis highlighted 6 clusters. The first cluster consisted in patients with no particular immune infiltrate and was annotated NA. The second cluster was annotated M0 due to its enrichment in M0 macrophages. The third cluster was annotated TLS (tertiary lymphoid structures as it was enriched in CD8^+^ T cells, T follicular helper cells (Tfh), naïve B cells and M2 macrophages, which we recently demonstrated to improve prognosis in cervical tumors (13). The fourth cluster was composed M1 macrophages and regulatory T cells (Tregs) and was annotated M1 MΦ-Tregs. The fifth cluster was enriched in M1 and M2 macrophages, and was therefore named M1/M2 MΦ. Finally, the sixth cluster was enriched in CD8^+^ T cells, which are known to improve prognosis, and was named CD8^+^ T cells (**Figure 1, A**) (21). These clusters were used to generate OS Kaplan-Meier curves. For readability, we pooled the patients from M1 MΦ-Tregs and M1/M2 MΦ under the M1/M2 MΦ name. Here, we showed that the clusters enriched in M0 and M1/M2 MΦ were of poor prognosis compared to those enriched in TLS or CD8^+^ T cells (**Figure 1, B**).

**Figure 1.**
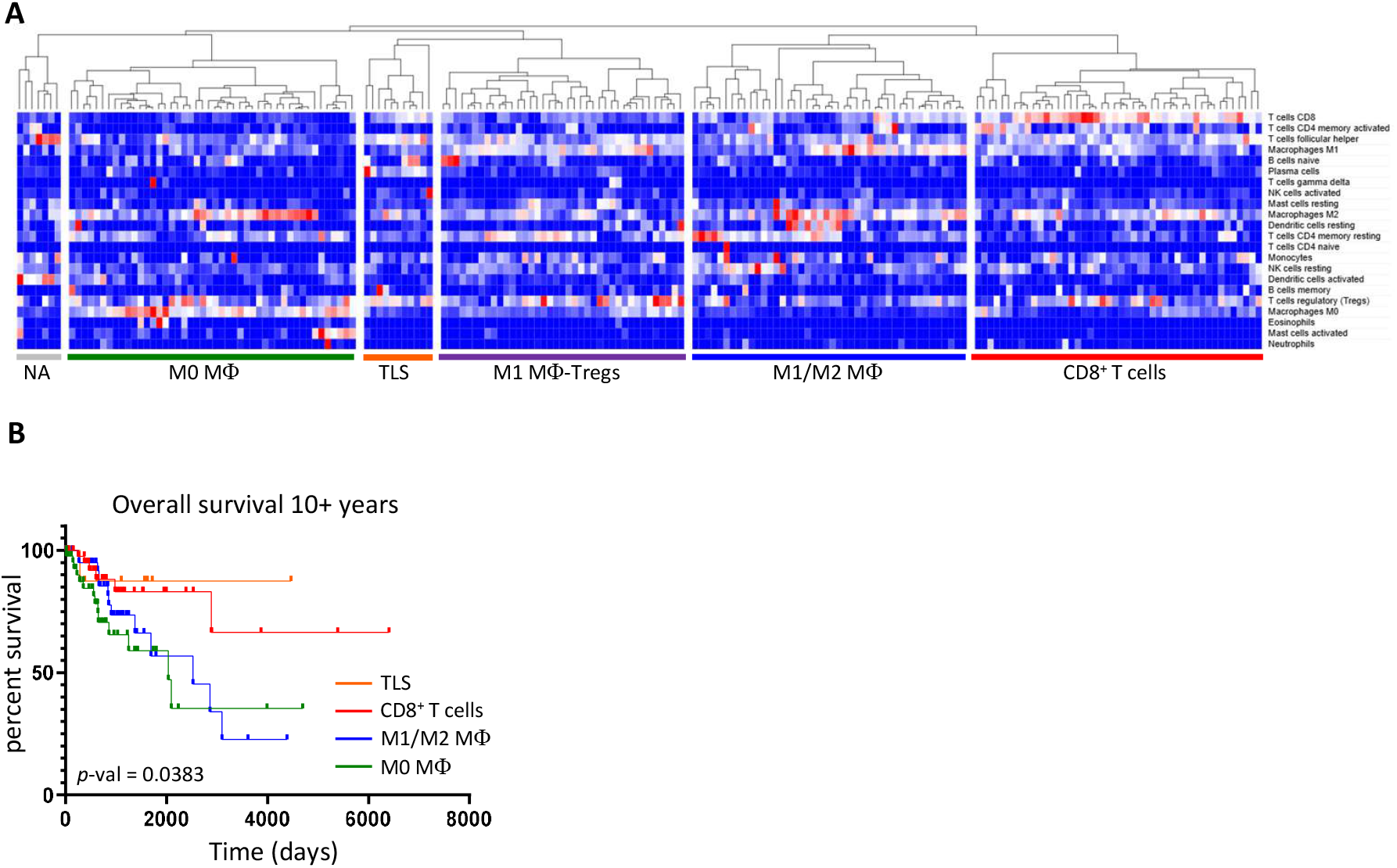
TAMs are of poor prognosis in cervical tumors Macrophages are markers of poor prognosis in cervical tumors. Publicly available TCGA RNAseq data were enriched for immune cells using the LM22 signature in CibersortX. Enrichment values were plotted in a heatmap and clustered according to cell types and samples using unsupervised hierarchical clustering. Main clusters were annotated according to immune cell enrichment determining the main TMEs in cervical tumors (**A**). Immune cell enrichment and TCGA patient vital status data (live/dead) were used to determine patient overall survival (OS), which was plotted on a Kaplan-Meier graph (**B**).

### TAMs and DCs express NECTIN-2 and PVR in cervical tumors

Cervical tumors are composed in majority of squamous cell carcinomas (SCC) the rest of tumors being divided between adenocarcinomas (ADK), mucinous and adenosquamous carcinomas. Here, we investigated the expression of PVRL receptor family in the CESC mRNAseq gene expression dataset from the TCGA. Log2 gene expression of PVR, Nectin-1, Nectin-2, Nectin-3 and Nectin-4 was extracted and compared between SCC, ADK, mucinous and adenosquamous cervical tumors (**Figure 2, A**). While the expression of each molecules remained overall high, some differences can be noted. PVR expression remained stable between the different types of tumors. Nectin-1 expression increased in SCC compared to ADK and mucinous carcinomas. Nectin-2 expression increased in mucinous carcinomas compared to SCC but not ADK or adenosquamous tumors. Nectin-3 increased in mucinous and ADK compared to SCCs, while Nectin-4 increased in SCC compared to mucinous and ADK (**Figure 2, A**).

**Figure 2.**
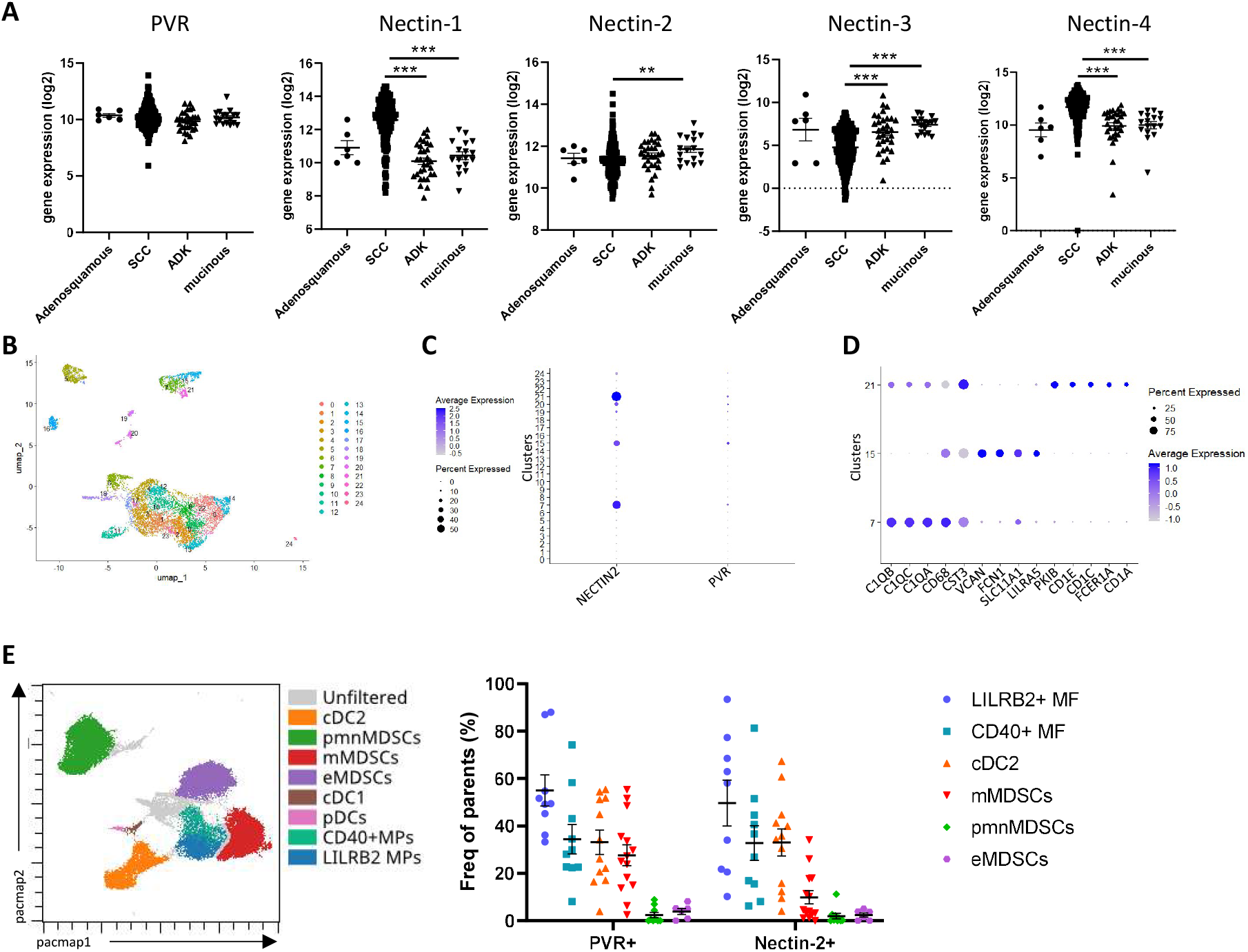
TAMs and DCs express NECTIN-2 and PVR in cervical tumors TAMs and cDC2 express NECTIN-2 and PVR in cervical tumors. TCGA dataset was used to assess the overall expression of PVR, NECTIN-1, NECTIN-2, NECTIN-3 and NECTIN-4 in cervical cancer (CESC) (**A**). Cervical cancer scRNA-seq datasets were downsampled on PTPRC^+^ (CD45^+^) immune cells and used to investigate cell-specific NECTIN-2 and PVR expression. UMAP projection and SNN-determined PTPRC^+^ immune cell clusters are shown in panel **B**. NECTIN-2 and PVR expression are displayed among SNN-identified cell clusters (**C**). The top 5 markers from DGE are shown for clusters 7, 15 and 21 allowing their identification (**D**). Mass cytometry was used to assess PVR and Nectin-2 expression in Tumor and tumor infiltrating myeloid cells from 15 cervical cancer samples. Gated cell phenotypes were projected onto a PaCMAP (left) and PVR and Nectin-2 frequency of expression was quantified (right) (**E**). Statistical significance was assessed by non-parametric 2-way ANOVA. \**p*-val<0.05, \*\**p*-val<0.01, \*\*\**p*-val<0.005.

To address the specificity of Necin-2 and PVR in the TME (3,5) we used scRNA-seq data from 4 different datasets (n=18 samples including 14 tumors and 4 healthy samples). Datasets were merged, subsampled for PTPRC+ (CD45+) cells, filtered and normalized as described in the methods section. We used the top 800 highly variable genes and the 20 first components of the PCA to generate a UMAP of cervical tumors infiltrating immune cells. Using SNN, we identified 25 clusters of immune cells (**Figure 2, B)**. Among these clusters, only clusters 7, 15 and 21 expressed Nectin-2 and PVR, although PVR expression was quite weak at the transcript level (**Figure 2, C**). Interestingly no statistical differences were found in the expression of PVR nor Nectin-2 when we compared their expression among clusters according to healthy, ADK or SCC samples (**Supp. Figure 1 A, B**). A differential gene expression (DGE) analysis was performed and the top 20 genes were used to identify cell types from the 25 clusters (**Supp. Table 2**). The top 5 gene expression from cluster 7, 15 and 21 were plotted. Cluster 7 had an increased expression of C1QA, B and C as well as CD68 and CST3, which allowed to identify this cluster as C1Q^+^ macrophages. Cluster 15 was enriched in cells expressing CD68, VCAN, FCN1, SLC11A1 and LILRA5, which allowed to annotate them as FCN1+ macrophages. Cluster 21 cells expressed higher levels of CST3 and CD1A, C and E as well as FCER1A which allowed to annotate them as conventional dendritic cells 2 (cDC2) (**Figure 2, D**). Using the top 20 differentially expressed genes in cluster 7, 15 and 21 we identified markers to validate our findings using markers existing in our mass cytometry panel. We identified CD1a and CD1c as markers for cluster 21 (cDC2), LILRB2 for cluster 7 (C1Q MΦ) and CD40 for cluster 15 (FCN1 MΦ).

Then, we investigated PVR and Nectin-2 expression in cervical tumors by mass cytometry (n=15). Our panel (**Supp. Table 3**) allows the identification of Macrophages subsets (CD33^+^CD11b^+^HLA-DR^+^CD68^+^CD64^+^LILRB2^+/-^CD40^+/-^), Monocytes (CD33^+^CD11b^+^HLA-DR^+^CD68^-^CCR2^+^CD14^+^), monocytic (m)MDSCs (CD33^+^CD11b^+^HLA-DR^lo/-^CD14^+^) and granulocytic (g)MDSCs (CD33^+^CD11b^+^HLA-DR^lo/-^CD15^+^) and cDC2 (HLA-DR^hi^CD11c^+^CD1a^-/+^CD1c^+^) (**Figure 2, E**). We found that LILRB2^+^ MΦ expressed PVR and Nectin-2 the most compared to other myeloid cell types (**Figure 2, E**). cDC2 and CD40^+^MΦ also expressed higher level of PVR and NEctin-2 compared to MDSCs. Interestingly, the expression of PVR, Nectin-2 or their co-expression did not vary according to tumor severity. Indeed, no differences could be observed between low risk (FIGO stage I + II) compared to high risk (FIGO stage III + IV) in Tumor cells, LILRB2^+^ MΦ, CD40^+^ MΦ macrophages and Monocytes (**Supp. Figure 1, C**). Similarly, we could not observe any significant difference of expression, or co-expression between SCC and ADK, although CD40^+^ MΦ and LILRB2^+^ MΦ macrophages tend increase Nectin-2 expression in SCC (**Supp. Figure 1, D**). Altogether, these data confirmed scRNA-seq data and identify TAMs as the major component of Nectin-2 and PVR mediated inhibition.

### Anti-PVR and Nectin-2 mAbs antagonistic mAbs reverse TIGIT -induced M2-like phenotype in MDMs

To investigate whether blocking PVR and Nectin-2 signaling would affect macrophage phenotypic profile, we generated monocyte-derived macrophages (MDM). MDM, when differentiated using MCSF, tend to spontaneously acquire immunosuppressive “M2” phenotype. Therefore, before the end of MDM differentiation, we added IFN-γ to polarize them towards an inflammatory “M1” phenotype (MDM1). We then gate differentiated MDM (**Figure 3, A**) and assessed PVR and Nectin-2 expression (**Figure 3, B**). MDM expressed PVR and Nectin-2 compared to unstained MDM and T cells, as shown in **Figure 3, B**. Then, we investigate MDM1 response to TIGITfc (soluble form of TIGIT). For that MDM1 were culture in the presence of TIGITfc (5µg/ml) and were according the following conditions, Isotype control (IgG1, 10µg/ml), anti-PVR (5µg/ml), anti-Nectin-2 (5µg/ml) and anti-PVR+anti-Nectin-2 combination (Combo, 10µg/ml). We performed flow cytometry in these conditions to identify phenotypic alterations in MDM1. As reported previously, TIGITfc induced the expression of immunosuppressive markers (CD163, CD209), which was not altered by the addition of IgG1 isotype. The expression of CD206, CD86, HLA-DR and CD80 did not significantly vary among the different conditions. However anti-Nectin-2 induced significant expression of CD83 compared to IgG1 isotype and other conditions. Anti-Nectin-2, and mainly the Combo condition induced the loss of CD163 and CD209 expression compared to IgG1 isotype (**Figure 3, C**). Here, we demonstrate that MDM can be used to assess anti-PVR and Nectin-2 efficiency and that TIGIT binding to its ligand induces an immunosuppressive profile. Finally, we show that anti-PVR and anti-Nectin-2 combinations allow to repolarize MDM2 towards inflammatory MDM1 profile as marked by the loss of CD209 and CD163 and the gain of CD83.

**Figure 3.**
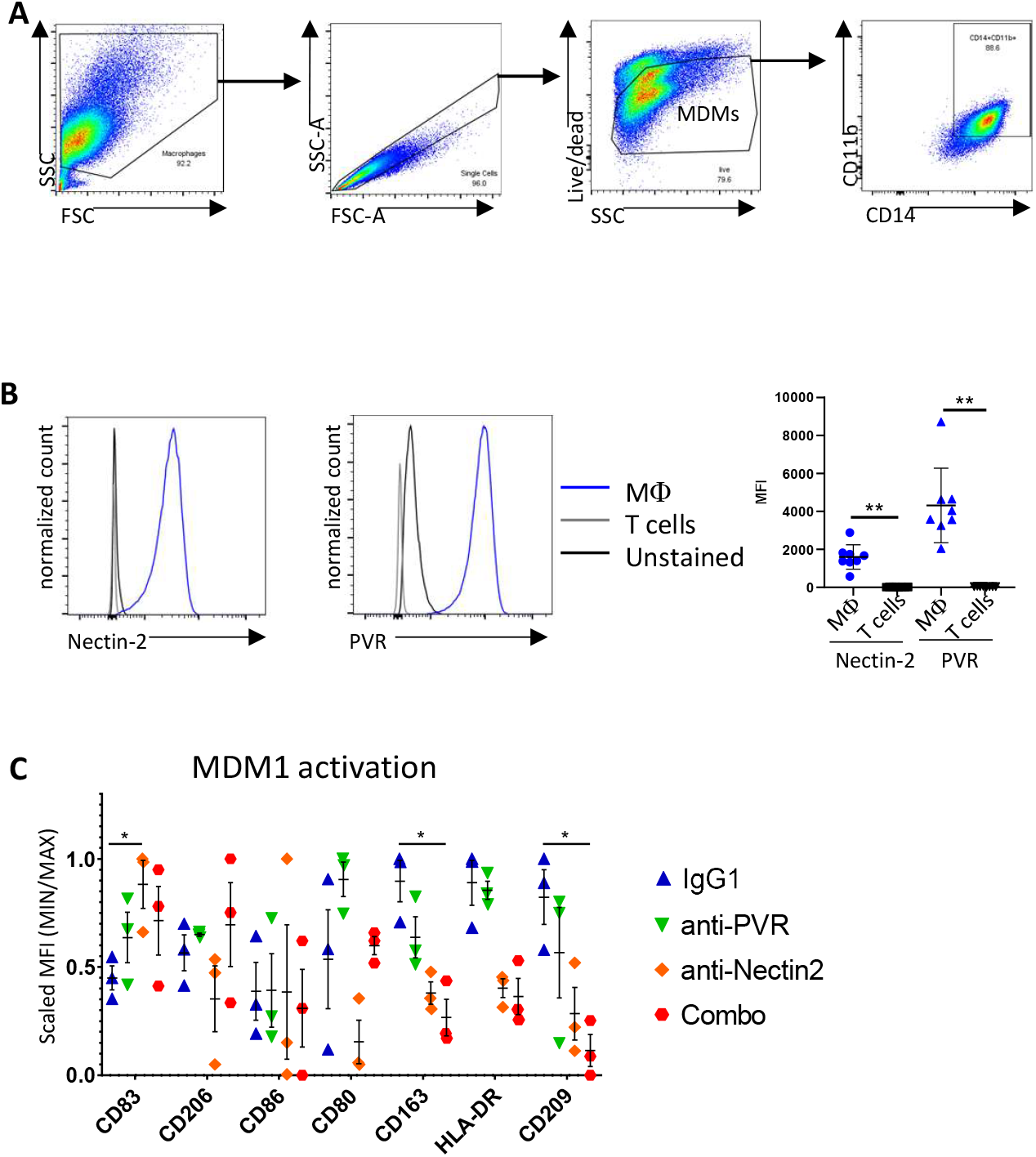
Anti-PVR and Nectin-2 mAbs antagonistic mAbs reverse TIGIT-induced M2-like phenotype in MDMs. Targeting PVR & Nectin 2 repolarize inhibitory macrophages towards inflammation. MDM1 were differentiated using MCSF and IFN-γ during the last 2 days. They were then exposed to TIGIT-fc to induce a immunosuppressive program in the presence or absence of control IgG1 (10µg/ml), anti-PVR (10µg/ml), anti-Nectin2 (10µg/ml) or Combo (10µg/ml). Cells were then stained for flow cytometry and gated (**A**). Nectin-2 and PVR expression by MDM were quantified by flow cytometry and compared to unstained MDM and T cells (**B**). Flow cytometry was performed to determine MDM activation state following the previously described conditions (**C**). Statistical significance was assessed by non-parametric 2-way ANOVA. \**p*-val<0.05.

### Anti-PVR mAbs restores phagocytosis of cervical tumor cell line in M2-like MDMs

Phagocytosis is a critical function of macrophages, which may participate to tumor elimination. Indeed, targeting “don’t eat me” signals (Magrolimab, anti-CD47) is an efficient strategy to control tumor growth via the activation of macrophages and other phagocytes. Here, we assessed the potential of anti-PVR and anti-Nectin-2 in restoring MDM phagocytosis. We used high-throughput confocal microscopy to track fluorescent cervical tumor cell line (mKate2-SiHa) phagocytosis by PKH67-labelled MDMs at different timepoints (30min, 60min, 120min and 240 min). We measured the area of mKate2 signal within a macrophage (area of red within green) and the percentage of phagocytosed SiHa as readouts. We observed that anti-PVR and Combo induced phagocytosis as early as 30min after the start of co-culture with SiHa, while Magrolimab-induced, our phagocytosis positive control, peaked at 60 min (**Figure 4**). This was observed in terms of area of SiHa engulfed into MDMs (area of red into green, **Figure 4, B**) and in terms of frequency of phagocytosed cells (**Figure 4, C**). Phagocytosis was maintained with anti-PVR up to 120min post co-culture, which was not the case for Magrolimab. After 60min, phagocytosis seemed to be reduced in Magrolimab and Combo conditions probably because of the diminution of SiHa available (**Supp. Figure 2, A, B, C**). Taken together, our results show that anti-PVR is a potent phagocytosis inducer and that combining it with anti-Nectin-2 does not alter this function.

**Figure 4.**
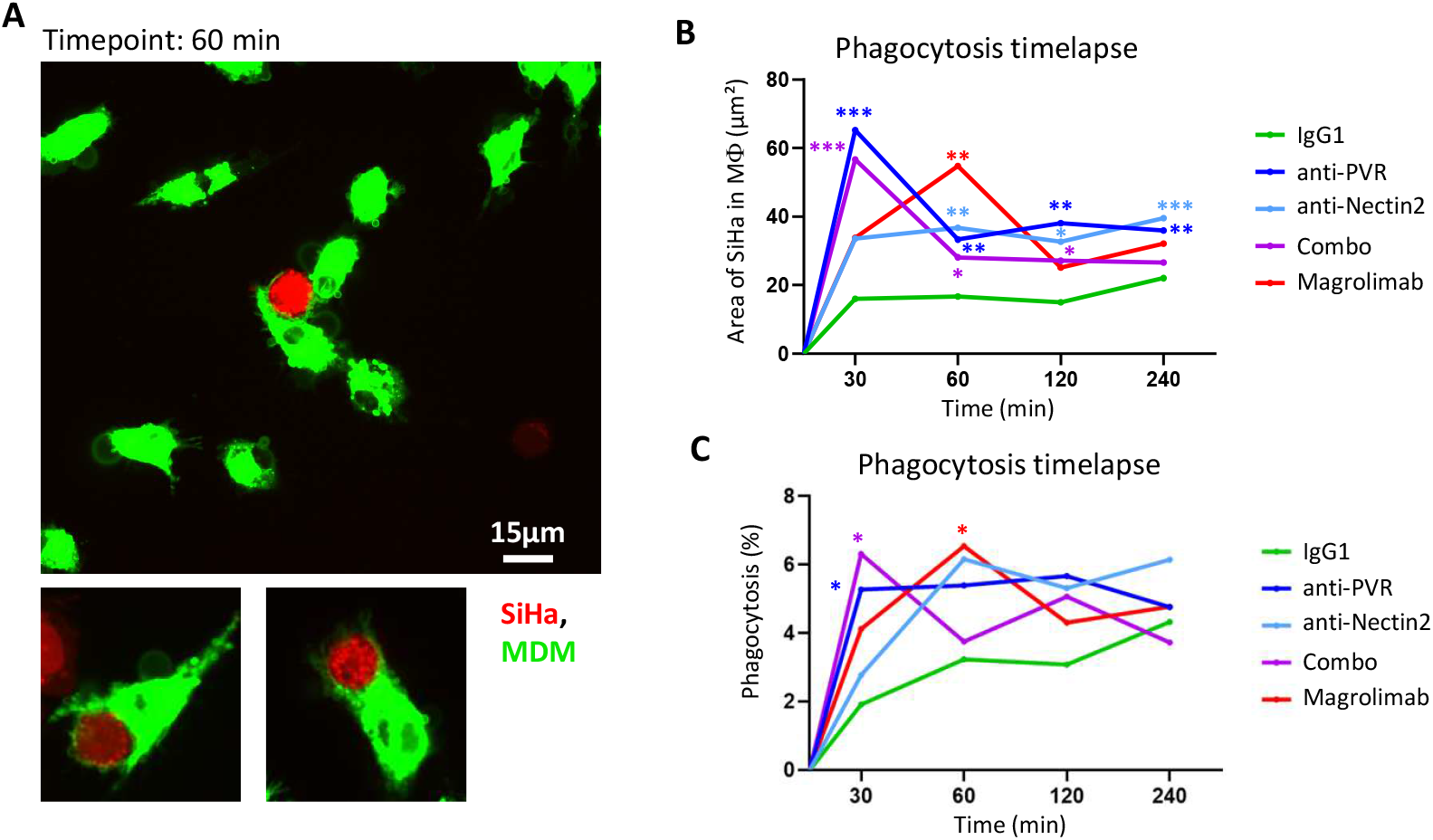
Anti-PVR mAbs restores phagocytosis of cervical tumor cell line in M2-like MDMs. Targeting PVR & Nectin-2 enables phagocytosis. MDM were differentiated as described and stained using PKH67. They were then cocultured with mKate2-SiHa cell line at a ratio of 1/5 for 30min, 60min, 120min and 240min in the presence of control IgG1 (10µg/ml), Magrolimab (10µg/ml), anti-PVR (10µg/ml), anti-Nectin-2 (10µg/ml) or both antibodies (Combo, 10µg/ml). Representative images of confocal microscopy are displayed (A) showing SiHa (red) being phagocytosed by MDM (green) (**A**). We calculated the area of SiHa present inside MDM vacuoles at 4 different timepoints and calculated the significance of phagocytosis compared to the control IgG1 stimulation (**B**). Phagocytosis frequency per condition and per timepoint was calculated and significance was assessed (**C**). P-values were calculated by non-parametric ANOVA by comparing values of other conditions to that of control IgG1. \**p*-val<0.05, \*\**p*-val<0.01, \*\*\**p*-val<0.005.

### Targeting PVR and Nectin-2 control tumor growth in both 2D and 3D cultures

Macrophages are known to participate to tumor invasion by remodeling the extracellular matrix (ECM) and secreting effectors such as growth factors EGF and TGF-β, which enhance tumor cell proliferation. Therefore, we hypothesized that immunosuppressive MDM (MDM2) would enhance SiHa proliferation. We performed a wound-healing assay were SiHa are cultured in chambers separated by a wall. The wall is removed and SiHa from each side of the wall migrate towards each other. We quantified the speed of gap-closing by measuring the frequency of empty-area between cells. We compared SiHa alone and SiHa with MDM2 and observed a faster gap-closing, hence proliferation, of SiHa in the presence of MDM2 (**Supp. Figure 3**). Next, we assessed the potential of anti-PVR and anti-Nectin-2 to dampen SiHa proliferation. Therefore, we performed a similar experiment including the following conditions, all in the presence of MDM2 and TIGITfc (n=5 MDM donors), isotype (IgG1, 10µg/ml), anti-PVR (5µg/ml), anti-Nectin-2 (5µg/ml) and anti-PVR + anti-Nectin-2 (Combo, 10µg/ml) (**Figure 5, A**). We found that only anti-PVR (green) slowed SiHa proliferation significantly, while anti-Nectin-2 and Combo did not show any significant difference. This is illustrated by the far-right graph comparing anti-PVR to IgG1 (**Figure 5, A**).

**Figure 5.**
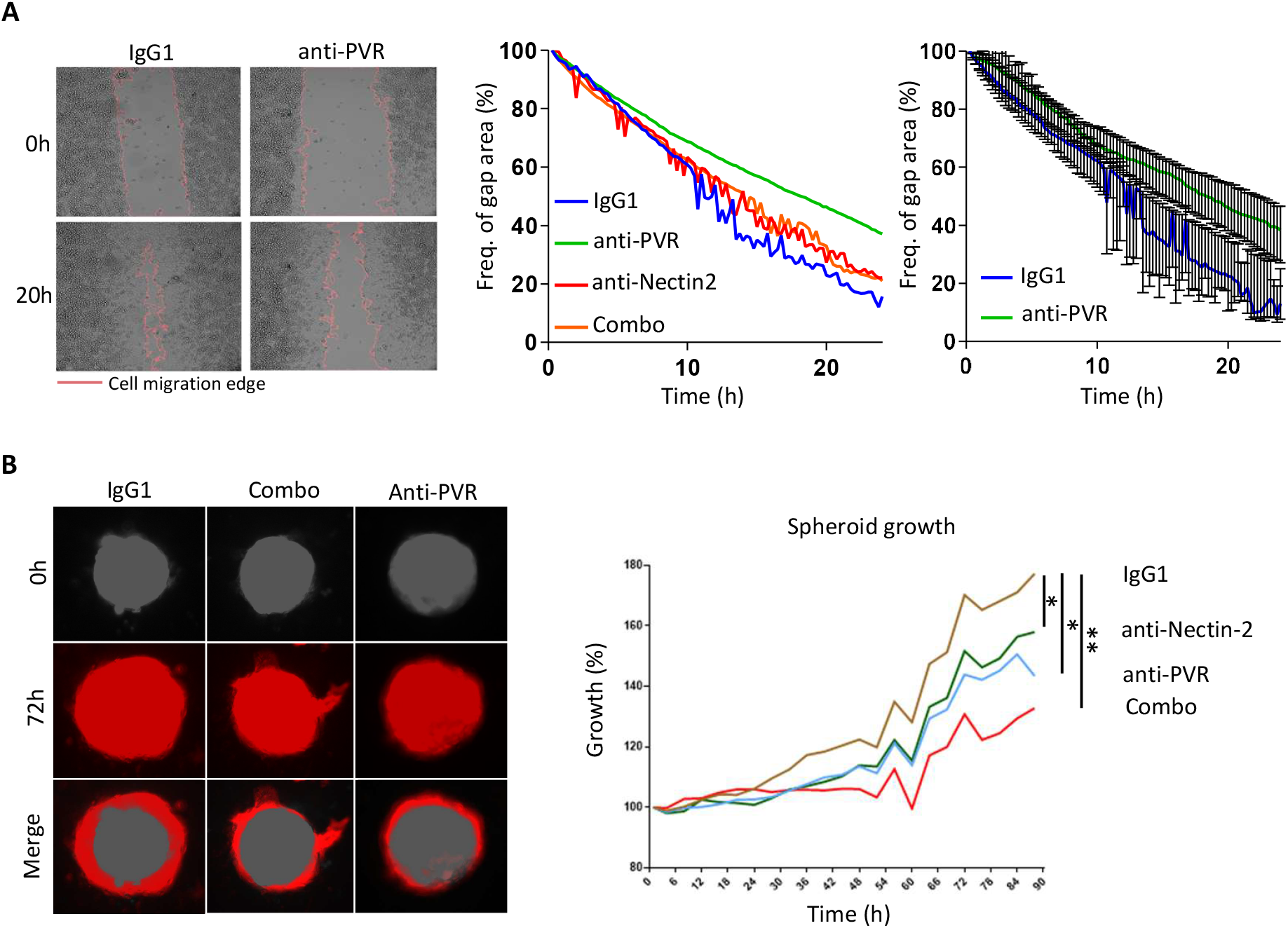
Targeting PVR and Nectin-2 control tumor growth in both 2D and 3D cultures. Targeting PVR & Nectin2 reduce tumor growth *in vitro*. MDM and SiHA were co-cultured in inserts composed of two separated chambers in the presence of TIGIT-fc in the presence or absence of control IgG1, anti-PVR, anti-Nectin2 or Combo. Inserts were then removed to let tumor cells migrate towards each other and fill the gap (**A, left**). Gap area was then quantified by video microscopy for 24h and plotted according to the different conditions (IgG1 blue, anti-PVR green, anti-Nectin2 red or Combo orange) (**A, center**). Anti-PVR showed a significant difference in slowing SiHa growth compared to control IgG1 (**A, right**). PVR and Nectin2 expressing MDM were differentiated and cocultured with SiHa to form spheroid during 5 days. Spheroids were then cultured with TIGIT^+^ T cells and control IgG1, anti-PVR, anti-Nectin-2 or Combo at 10µg/ml. Spheroid growth was assessed by video microscopy (**B**). P-values were calculated by non-parametric ANOVA by comparing values of other conditions to that of control IgG1. \**p*-val<0.05, \*\**p*-val<0.01.

Knowing that anti-PVR and Nectin-2 add effect on macrophage polarization and in controlling tumor cell proliferation, we built a spheroid culture system of immune infiltration. For this experiment we chose the mKate2-Caski fluorescent cervical cancer cell line, which spontaneously forms spheroids in low adherence conditions. The spheroids were formed using mKate2-Caski and MDM2 during 5 days. In the meantime, autologous T cells were stimulated using anti-CD3/CD28 beads during 3 days to allow the expression of TIGIT at their surface. Finally, we cocultured the spheroids with activated T cells during 90h and imaged spheroid growth by video-microscopy in the following conditions: isotype (IgG1, 10µg/ml), anti-PVR (5µg/ml), anti-Nectin-2 (5µg/ml) and anti-PVR + anti-Nectin-2 (Combo, 10µg/ml) (**Figure 5, B, left**). We observed that anti-PVR and anti-Nectin-2 were able to slow tumor growth by 50% at 90h compared to IgG1 (**Figure 5, B, right**). The Combo condition reached 70% of growth reduction at 90h compared to the IgG1. Taken together our results show that blocking PVR/Nectin-2-TIGIT interaction allows tumor growth control in vitro in 2D or in more complex 3D models.

### Targeting PVR and Nectin-2 restores anti-tumor response in dynamic cervical tumor 3D cultures

Considering the results on tumor size control, we investigated the potential of our treatment to recruit immune effectors following treatment. Briefly, patient tumor samples were cultured into 25% matrigel on a 8µm transwell filter overnight in the presence of IgG1, anti-PVR, anti-Nectin-2, both antibodies (Combo) or Atezolizumab, all at a concentration of 10µg/ml. We used Atezolizumab as this mAb was previously used in the Skyscraper trial and improved patient survival. Furthermore, Atezolizumab targets PD-L1 which may also be expressed by TAMs and the filter containing tumor sample culture was then put on the MIVO system as described in the method section. Autologous CD45-APC stained PBMCs were loaded into the millifluidics and allowed to flow in the system overnight. Transwell filters were then removed, dissociated and stained for spectral cytometry (**Figure 6, A**). We then measured the frequency of PBMCs (CD45-APC+) infiltrating the tumor compartment. To verify the physiological behavior of our assay, we first assessed the potential of hot tumors to recruit more immune cells from the circulation compared to cold tumors. We used the presence or absence of tertiary lymphoid structures (TLS) as a marker of hot (TLS^+^) or cold (TLS^-^) tumors. We showed that TLS+ tumors recruited more immune cells than TLS negative tumors. Indeed, the recruitment of all immune cell types (γδ T cells, CD4^+^ T cells, CD8^+^ T cells, B cells, Myeloid cells and NK cells) was more intense in the experiments with TLS^+^ tumors. As hot tumors are known to be enriched in recruited immune cells, we validated our recruitment model. Next, we focused on the response to immunotherapies. Here, we demonstrated that, in comparison to control IgG1 treatment, some samples exhibited an increase in PBMCs recruitment and others did not show differences or showed a reduction in recruitment (**Figure 6, B**). Interestingly, Patient #1, #2, #4 and #7 increased PBMC recruitment following Atezolizumab stimulation, which was different with other treatments. Indeed, anti-PVR treatment only increased recruitment in patient #5, anti-Nectin-2 increased recruitment in patients #1, #3 and #5 and combination treatment (Combo) in patients #1, #3 and #5. We concluded that patients #1 was susceptible to any of our treatments, while patients #2 and #4 were only susceptible to Atezolizumab, and patients #3 and #5 were rather affected by anti-PVR, anti-Nectin-2 or the combination (**Figure 6, C**). Taken together our results show an increase in autologous PBMCs recruitment following treatment. This recruitment seems dependent on the sensitivity of tumors to the blockade of PD-L1 or PVRL.

**Figure 6.**
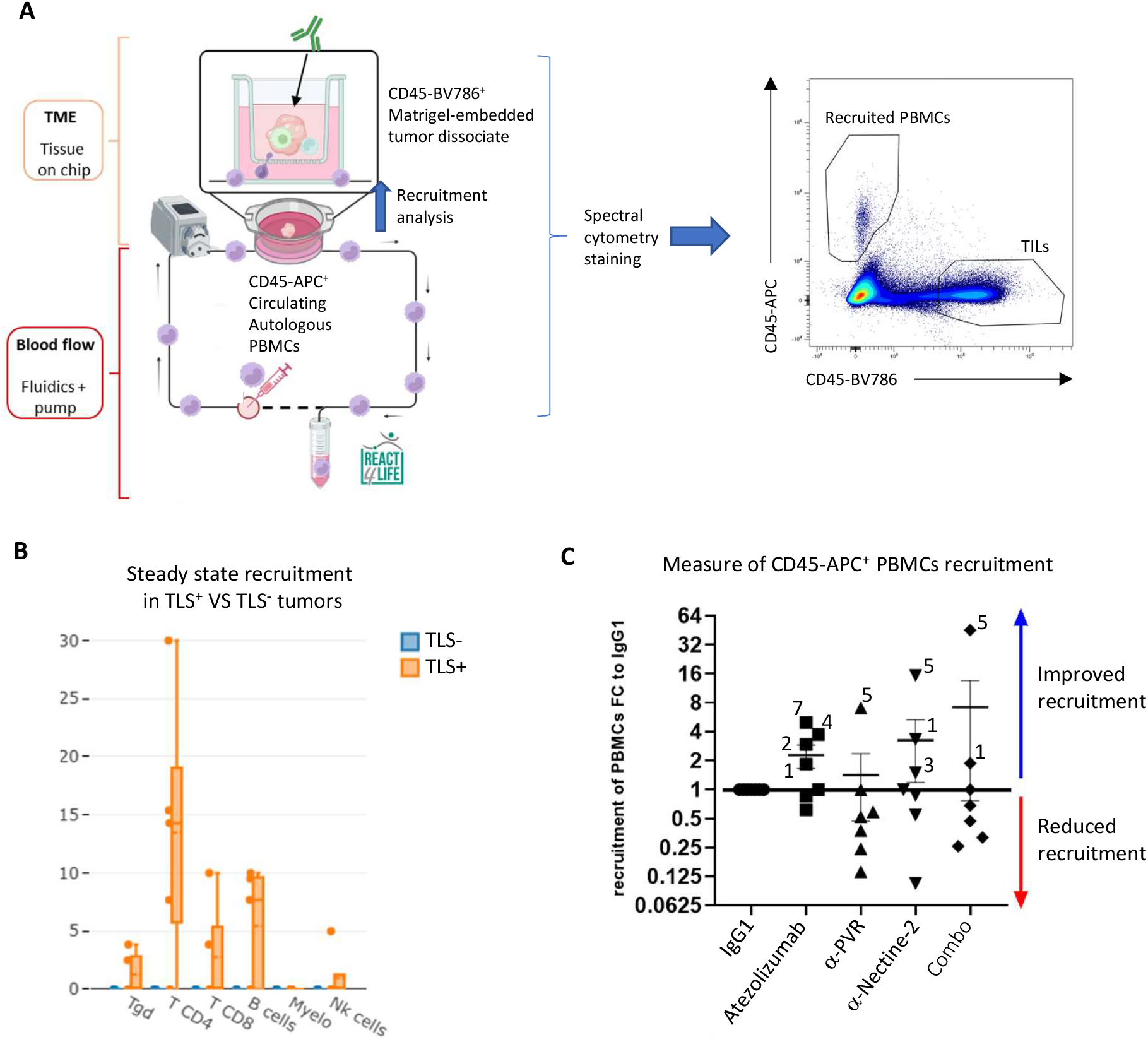
Targeting PVR and Nectin-2 restores anti-tumor response in dynamic cervical tumor 3D cultures. Targeting PVR & Nectin-2 *ex vivo* increases immune cell recruitment in a millifluidic 3D culture system. Patient tumor samples (n=7) were thawed, rested stained for CD45-BV786 and cultured into matrigel on 8µm transwell filter overnight in the presence of IgG1, anti-PVR, anti-Nectin-2, both antibodies (Combo) or Atezolizumab. The filter containing tumor sample culture was then put on the MIVO system as described in the method section. Autologous CD45-APC stained PBMCs were loaded into the millifluidics. Peristaltic pump was then turned on letting PBMCs flow in the system overnight. Transwell filters were then removed, dissociated and stained for spectral cytometry. Filter supernatants were saved for Luminex dosages (**A**). Cell infiltration was determined at steady-state (treatment free) for TLS+ and TLS-tumors. We assessed the frequency of infiltrating gd Tcells, CD4^+^ T cells, CD8^+^ T cells, B cells, Myeloid cells and NK cells among total infiltrate (**B**). Cell migration was assessed by spectral cytometry in the dissociated cell from the transwell filter by CD45-APC^+^ gating and compared to that generated by IgG1 isotype control. When the frequency of infiltrating cells was superior to that of IgG1, patients were considered as “responders” (blue arrow) and those with equal or inferior frequency were considered as “non-responders” (red arrow) (**C**).

## Discussion

The investigation of TAM response to ICB is an expending field aiming at enhancing response to conventional ICBs such as pembrolizumab or atezolizumab (1). Indeed, TAMs are known to participate to tumor escape from ICB or conventional treatments (22–24). Among other methods such as designing chimeric antigen receptor Macrophages (CAR-M) (25) or small molecules, the use of mAbs directly targeting TAMs or aiming at restoring their function represent one of the main strategies to restore anti-tumor immunity. Indeed, “don’t eat me signal” targeting ICBs to restore tumor cell killing by phagocytosis are in development or reached clinical trial (Magrolimab for example) (26–29). Another strategy is to force myeloid cell or TAM activation by using agonists of receptors such as CD40 (30) or even Toll-like receptors ligands (31,32). In our study we focus on the myeloid side of the TIGIT-PVRL axis. Anti-TIGIT mAbs such as Tiragolumab modestly improved pembrolizumab efficiency. As mentioned, a study demonstrated that using ADCP-triggering mAbs instead of IgG4 would improve treatment by stimulating myeloid cells. We decided to target TIGIT ligands PVR and Nectin-2, which are both expressed by TAMs in cervical tumors. Our approach aimed at repolarizing TAMs towards inflammatory profiles to trigger adaptive immunity tumor cell killing. PVR was known to induce a regulatory signal in DCs via the ERK pathway and the release of IL-10 (33). We showed that blocking PVR slowed cervical tumor cell line growth in coculture with MDM in 2D and in 3D. PVR was also described as a secondary “don’t eat me signal” participating to tumor escape from macrophage cytotoxicity (3). We could validate this aspect in cervical cancer models, as blocking PVR restored phagocytosis of the SiHa tumor cell line. Since PVR is not the only ligand of TIGIT present at the surface of TAMs, we anticipated that blocking solely PVR would only partly restore anti-tumor immunity. Indeed, anti-PVR treatment did not restore an inflammatory profile of MDMs, while Nectin-2 induced a decrease in the expression of anti-inflammatory/immunosuppression canonical markers CD163 and CD206, while restoring the expression of CD83, an Ig superfamily molecule shown to participate in antigen presentation and myeloid cell activation (14,34). However, blocking Nectin-2 did not affect SiHa phagocytosis as much as PVR-blockade, which suggests different pathway regulated by PVR and Nectin-2. Unfortunately, while PVR signaling is described and goes through the ERK/MAPK pathway, the signaling downstream of Nectin-2 is less studied and does not allow clear identification of the pathways affected by its blockade. The combination of both anti-PVR and Nectin-2 mAbs restored anti-tumor immunity in our model of spheroid growth. This suggest a total blockade of the interaction with TIGIT allowing the reinvigoration of TIGIT^+^CD8^+^ T cells, which therefore are able to prevent spheroid growth. Our study demonstrates the potential of a dual anti-PVR/Nectin-2 blockade in cervical cancer. *In vitro*, we could restore anti-tumor immunity in complex spheroid models. *Ex vivo*, in a complex model of millifluidics and 3D culture we could show an increase of immune cell recruitment following anti-PVR/Nectin-2 treatments. Interestingly, only a few clinical trials were interested in PVR blockade, and even less in Nectin-2 blockade (3). Interestingly, we also found that Atezolizumab induced immune cell recruitment in the same model, which was consistent with previous studies showing immune cell response to PD-L1 blockade. While our study is limited by the number of clinical samples used for functional assays, it paves the way for further studies using human modelization of the TME and associated immune cell recruitment. Indeed, TAM targeting treatment could be used in combination therapies prior or concomitantly to T cell targeting therapies, or to prepare TME to increase the efficiency of CAR T cells. With similar micro or millifluidics models, studies have shown efficiency of anti-PD-1 treatment, circulating tumor cell quantification and metastasis onset (35–38).

To conclude, our study shows for the first time that a dual blockade of PVR and Nectin-2 restores macrophages inflammatory functions and is efficient to reduce cervical tumor growth *in vitro* and *ex vivo* in pseudo-physiological models of autologous tumor cells, tumor infiltrating immune cells and PBMCs. These results may represent a proof of concept for targeting the TIGIT-PVRL axis on the myeloid cell side, and contribute to the understanding of macrophages response to immunotherapies.

## Supporting information

Supp.Table3_Spectral cytometry panel

Supp.Table1_Cibersort results

Supp.Table2_CyTOF panel

## Authors contribution

LG and OD conceptualized the project. LG, OD, NB, LGa performed formal analysis. LG, DO, MSD supervised the project. OD, NB, TF investigation. LG, NB, OD, ML, MSR, ABA, MR methodology, LG, OD, NB wrote the original draft. LG, DO, ASC and JAN edited the manuscript. LG, DO, AOB obtained resources and funding. RS, EL, XC performed data curation. LG, DO validated the project and manuscript. LG, DO and ASC were project administrators

## Acknowledgements

We would like to thank the cytometry platform (M. Dalmasso, A.L. Bailly), the microscopy platform (M. Rodrigues) and F. Bard team at CRCM for support and access to technologies. We would like to thank React-4-Life (S. Scaglione) for collaboration and advice for the use of the MIVO technology.

