## Supplementary material for "PVR and Nectin-2 blockade trigger macrophage anti-tumor functions, promote immune cell recruitment and prevent cervical tumor growth": Supp.Table3_Spectral cytometry panel

| **Excitation** | **Marker** | **Fluorochrome** | **Target specie** | **Manufacturer** | **RRIDs** |
| --- | --- | --- | --- | --- | --- |
| UV | CD11c | BUV395 | Human | Becton-Dickinson | AB_2872717 |
|  | CD16 | BUV496 | Human | Becton-Dickinson | AB_2744294 |
|  | CD4 | BUV805 | Human | Becton-Dickinson | AB_3684858 |
| Violet | IgD | BV421 | Human | Biolegend | AB_2561619 |
|  | CD38 | V450 | Human | Becton-Dickinson | AB_1937282 |
|  | CD56 | BV570 | Human | Biolegend | AB_2565918 |
|  | TCRgamma delta | BV605 | Human | Becton-Dickinson | AB_2742796 |
|  | CD127 | BV650 | Human | Biolegend | AB_2562095 |
|  | CD45 | BV785 | Human | Biolegend | AB_2563129 |
| Blue | HLA-DR | FITC | Human | Biolegend | AB_2563164 |
|  | CD3 | PerCP | Human | Becton-Dickinson | AB_10641841 |
|  | CD11b | PerCP Cy5.5 | Human | Becton-Dickinson | AB_394002 |
| Yellow/Green | CD25 | PE | Human | Biolegend | AB_2564145 |
|  | CD163 | PE-Dazzle594 | Human | Biolegend | AB_2890708 |
|  | TIGIT | PE-Cy7 | Human | Biolegend | AB_2632929 |
|  | CD33 | PE-Fire810 | Human | Biolegend | AB_2936534 |
| Red | CD45 | APC | Human | Biolegend | AB_2650648 |
|  | CD19 | SparkNIR685 | Human | Biolegend | AB_2860769 |
|  | CD8 | Alexa Fluor700 | Human | Becton-Dickinson | AB_3696551 |
|  | CD14 | APC-H7 | Human | Becton-Dickinson | AB_1645465 |
|  | CD27 | APC-Fire810 | Human | Biolegend | AB_2860962 |
|  | Live/dead | ViaDye Red | NA | CyTEK | NA |

**Supplementary Table 3:** Spectral cytometry panel.
