## Supplementary figures and images for "PVR and Nectin-2 blockade trigger macrophage anti-tumor functions, promote immune cell recruitment and prevent cervical tumor growth"

### Supp.Table1_Cibersort results

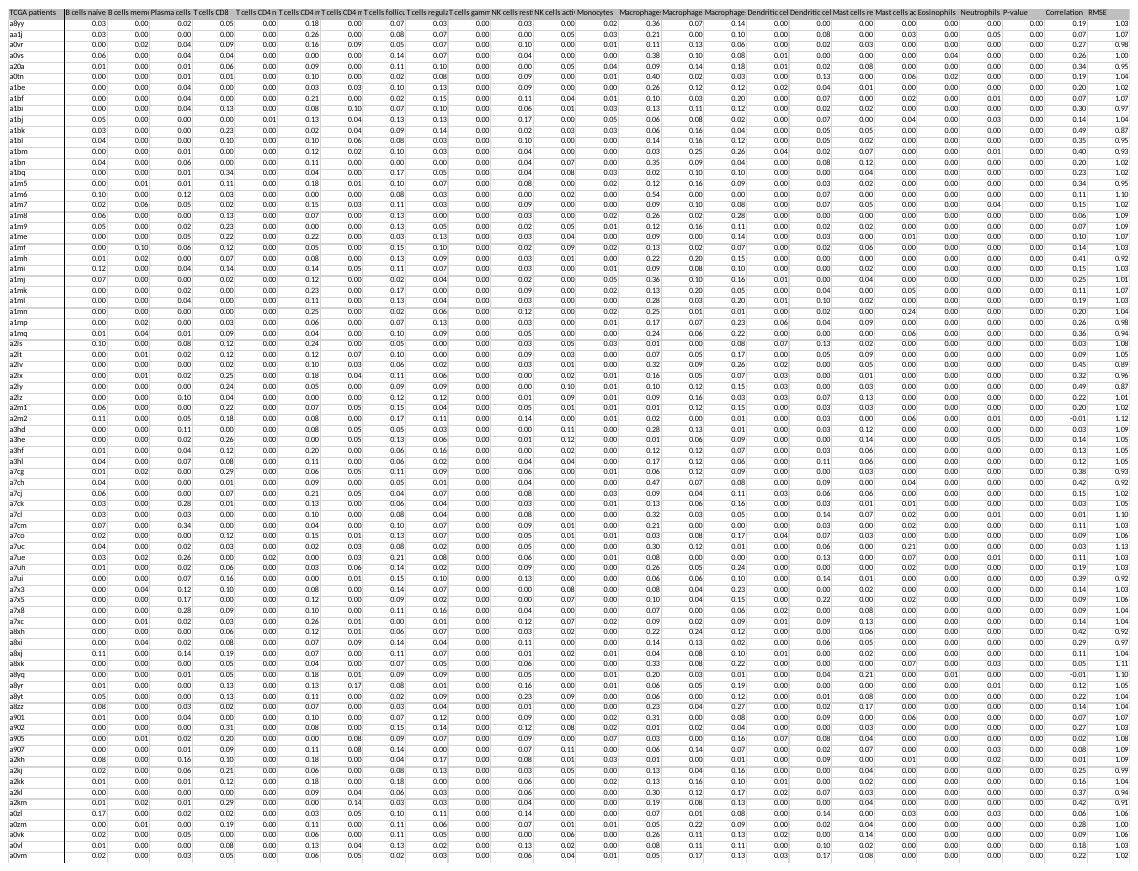

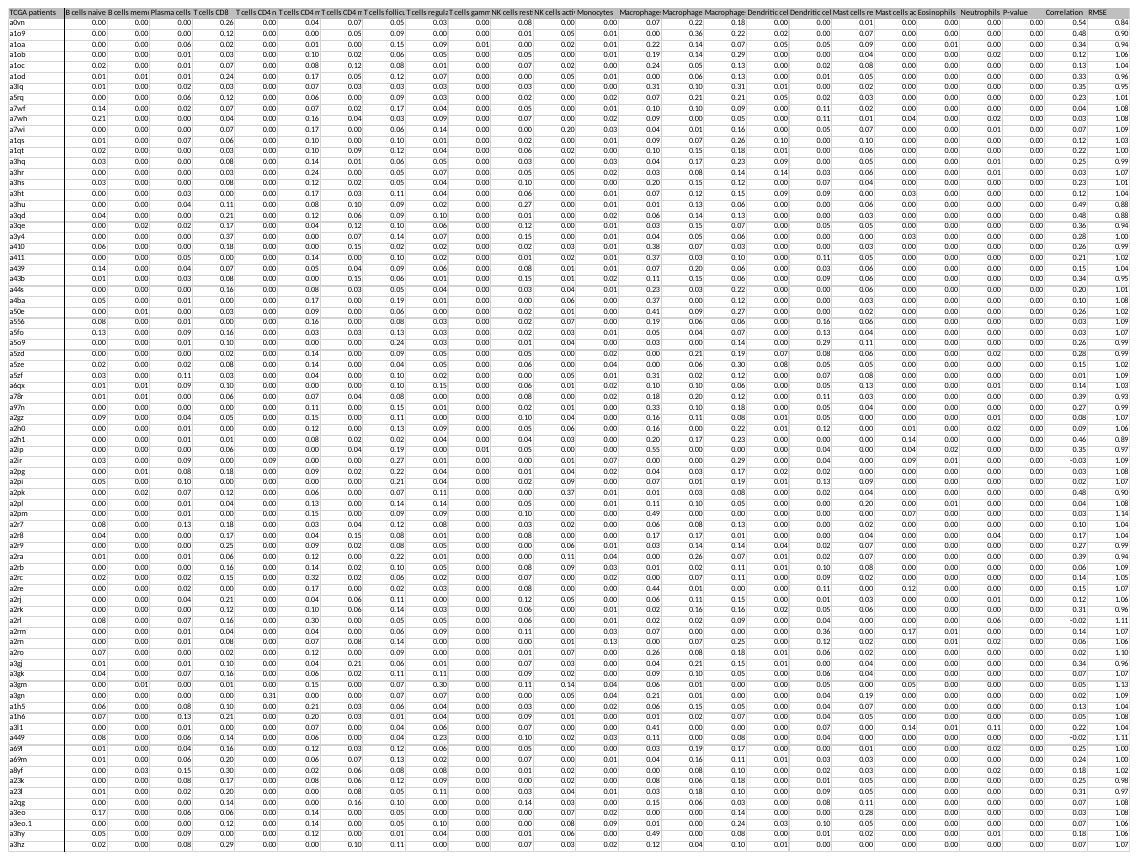

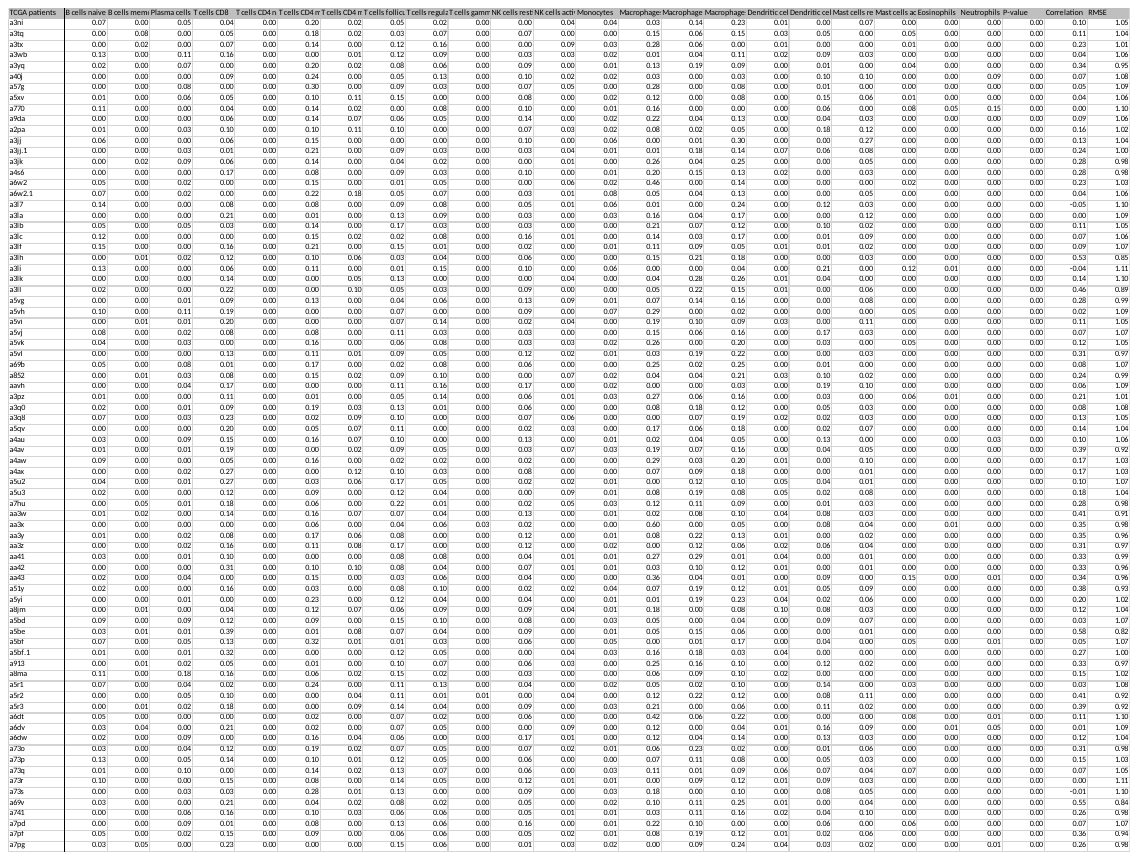

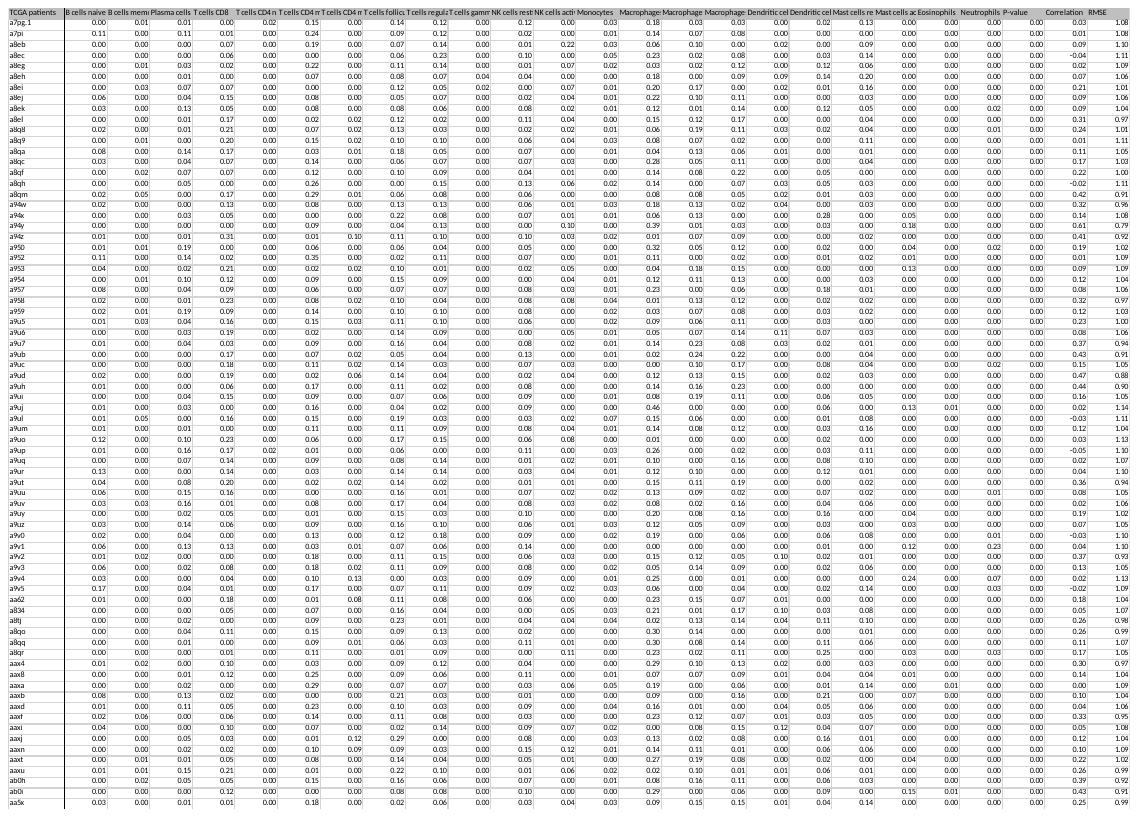
