## Supplemental Figures 1-2 for "PVR and Nectin-2 blockade trigger macrophage anti-tumor functions, promote immune cell recruitment and prevent cervical tumor growth"

### Slide 1
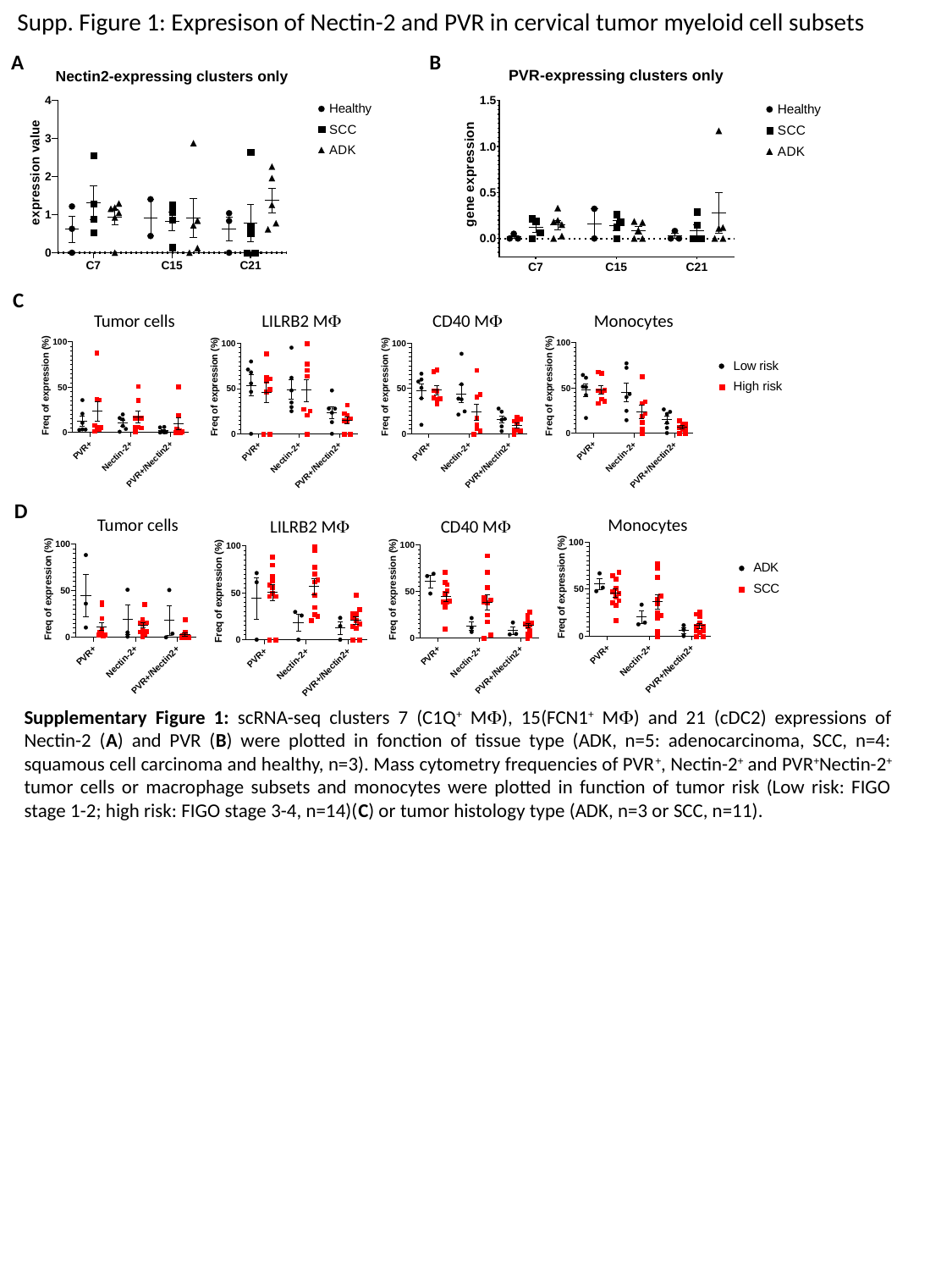

Supp. Figure 1: Expresison of Nectin-2 and PVR in cervical tumor myeloid cell subsets
A
B
C
Tumor cells
CD40 MΦ
Monocytes
LILRB2 MΦ
D
Tumor cells
Monocytes
CD40 MΦ
LILRB2 MΦ
Supplementary Figure 1: scRNA-seq clusters 7 (C1Q^+^ MΦ), 15(FCN1^+^ MΦ) and 21 (cDC2) expressions of Nectin-2 (A) and PVR (B) were plotted in fonction of tissue type (ADK, n=5: adenocarcinoma, SCC, n=4: squamous cell carcinoma and healthy, n=3). Mass cytometry frequencies of PVR^+^, Nectin-2^+^ and PVR^+^Nectin-2^+^ tumor cells or macrophage subsets and monocytes were plotted in function of tumor risk (Low risk: FIGO stage 1-2; high risk: FIGO stage 3-4, n=14)(C) or tumor histology type (ADK, n=3 or SCC, n=11).

### Slide 2
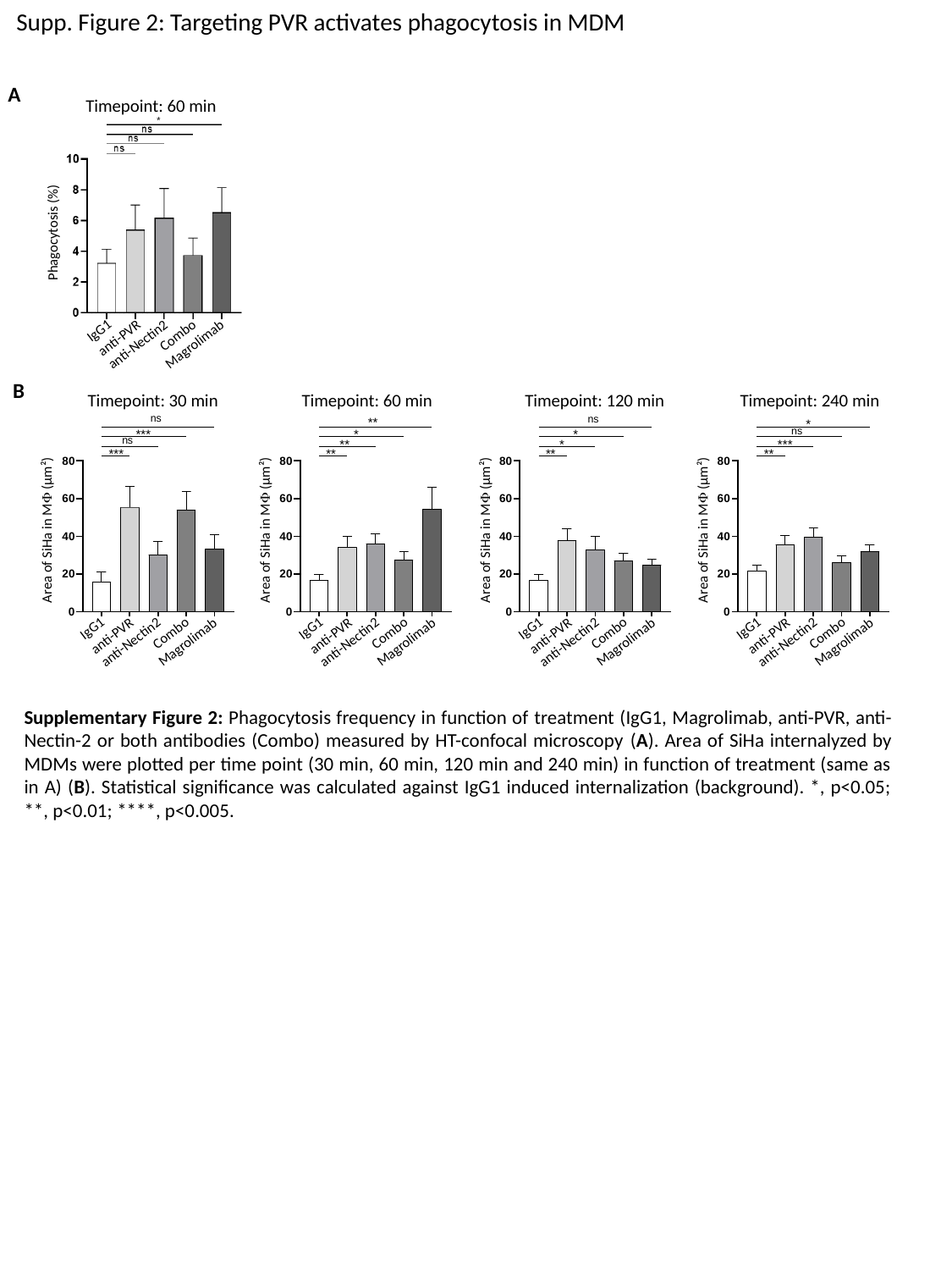

Supp. Figure 2: Targeting PVR activates phagocytosis in MDM
A
Timepoint: 60 min
Phagocytosis (%)
IgG1
Combo
anti-PVR
Magrolimab
anti-Nectin2
B
Timepoint: 30 min
Timepoint: 60 min
Timepoint: 120 min
Timepoint: 240 min
Area of SiHa in MΦ (µm²)
Area of SiHa in MΦ (µm²)
Area of SiHa in MΦ (µm²)
Area of SiHa in MΦ (µm²)
IgG1
Combo
anti-PVR
Magrolimab
anti-Nectin2
IgG1
Combo
anti-PVR
Magrolimab
anti-Nectin2
IgG1
Combo
anti-PVR
Magrolimab
anti-Nectin2
IgG1
Combo
anti-PVR
Magrolimab
anti-Nectin2
Supplementary Figure 2: Phagocytosis frequency in function of treatment (IgG1, Magrolimab, anti-PVR, anti-Nectin-2 or both antibodies (Combo) measured by HT-confocal microscopy (A). Area of SiHa internalyzed by MDMs were plotted per time point (30 min, 60 min, 120 min and 240 min) in function of treatment (same as in A) (B). Statistical significance was calculated against IgG1 induced internalization (background). *, p<0.05; **, p<0.01; ****, p<0.005.
